# Chick cranial neural crest cells migrate by progressively refining the polarity of their protrusions

**DOI:** 10.1101/180299

**Authors:** Miriam A. Genuth, Christopher D.C. Allen, Takashi Mikawa, Orion D. Weiner

## Abstract

*In vivo* quantitative imaging reveals that chick cranial neural crest cells throughout the migratory stream are morphologically polarized and migrate by progressively refining the polarity of their protrusions.

**Abstract:** To move directionally, cells can bias the generation of protrusions or select among randomly generated protrusions. Here we use 3D two-photon imaging of chick branchial arch 2 directed neural crest cells to probe how these mechanisms contribute to directed movement, whether a subset or the majority of cells polarize during movement, and how the different classes of protrusions relate to one another. We find that cells throughout the stream are morphologically polarized along the direction of overall stream movement and that there is a progressive sharpening of the morphological polarity program. Neural crest cells have weak spatial biases in filopodia generation and lifetime. Local bursts of filopodial generation precede the generation of larger protrusions. These larger protrusions are more spatially biased than the filopodia, and the subset of protrusions that power motility are the most polarized of all. Orientation rather than position is the best correlate of the protrusions that are selected for cell movement. This progressive polarity refinement strategy may enable neural crest cells to efficiently explore their environment and migrate accurately in the face of noisy guidance cues.

## Introduction

Directed cell migration is important for embryonic development, immune function, and wound healing. There are a number of possible strategies that cells can use to generate directed migration. They can read gradients of guidance cues spatially across their surface (Lauffenburger et al., 1987; Schwarz et al., 2017) and/or compare the concentration of agonist in time (Macnab and Koshland, 1972). Cells can bias the generation of protrusions or select among randomly-generated protrusions. Determining the logical framework cells use for migration is often challenging as it is not readily deduced from knockout phenotypes. However, careful quantitative imaging of migrating cells behavior can be used to discriminate between these migration strategies.

Quantitative imaging has been used to examine the guidance strategies in other migratory cells. Bacteria interpret gradients temporally and accomplish chemotaxis by increasing their run lengths that are directed up the chemoattractant gradient (Berg and Brown, 1972). Eukaryotic cells are capable of interpreting gradients spatially and can spatially bias the generation of protrusions for strong gradients (Arrieumerlou and Meyer, 2005; Zigmond, 1974). However, in more shallow gradients, cells show more random protrusion formation and instead spatially bias the selection of protrusions to accomplish directional movement (Andrew and Insall, 2007; Arrieumerlou and Meyer, 2005).

We have some understanding of the molecular logic linking protrusion dynamics to directed migration in Xenopus neural crest. These cells make use of a contact inhibition of locomotion (CIL) scheme for their migration (Carmona-Fontaine et al., 2008; Theveneau et al., 2010). CIL is a process in which a cell ceases forward movement upon contact with another cell (Abercrombie and Heaysman, 1953). In the case of sparsely distributed and weakly adhesive cells, CIL often induces the cell to reverse its polarity and move away from the site of contact.

However, if the cells are in a very crowded environment, as is the case of Xenopus neural crest cells, and/or are strongly adhesive they may be unable to disengage and will instead maintain contact even after suppression of protrusion formation at the site of contact. In Xenopus neural crest, cell-cell contact leads to cadherin engagement and activation of planar cell polarity (PCP) signaling to induce protrusion retraction at the site of contact (Carmona-Fontaine et al., 2008). As a result of this process, neural crest cells in the middle of the stream are non-polar and non-protrusive while those at the edge of the stream polarize and protrude. These multicellular contacts and PCP activation are necessary for the directed migration of Xenopus neural crest cells. In this setting, guidance cues act by modulating the lifetime of CIL-generated protrusions rather than inducing protrusions *de novo* (Theveneau et al., 2010).

To what extent the Xenopus model applies to cranial neural crest cell migration in amniotes is unclear. PCP signaling is required for neural crest migration in Xenopus but dispensable for neural crest cell migration in mice (Pryor et al., 2014). While the specific molecular mechanism of Xenopus CIL appears not to be conserved in mice, it does not exclude the possibility that the Xenopus overall strategy for neural crest migration could still apply.

Additionally, it has been reported that chick neural crest cells in the middle of the stream commonly have a bipolar morphology (Teddy and Kulesa, 2004), as opposed to the non-polar morphology of Xenopus neural crest, suggesting that chick may use a different mechanism than CIL to control overall cell shape.

To further explore the protrusion dynamics underlying neural crest cell migration we turned to chick as a model capable of live, *in vivo* imaging of migrating neural crest cells (Kulesa and Fraser, 1998). We find that neural crest cells in all regions of the migratory stream have polarized protrusion dynamics relative to the direction of overall stream movement. Neural crest cells show weak spatial biases in filopodia generation and lifetime, stronger biases in the generation of larger protrusions, and are most polarized in their productive protrusions. Thus, chick neural crest cells migrate by conducting a biased search with polarity refinement.

## Results

Because of the dense packing of neural crest cells during their migration in chick, it is difficulty to assay cell morphology when all cells are labeled. To analyze the distribution and dynamics of protrusions in this setting, we labeled a subset of cells in the stream by sparsely electroporating Hamburger and Hamiltion (HH) stage 8+ to 9 embryos with pCAGGS-Gap43GFP (mem-GFP to visualize the plasma membrane) and pCAGGS-H2B-tdTomato (to visualize the nuclei) and performed live 3D imaging at HH 11-12 via two-photon microscopy (Fig 1A). We found that neural crest cells in the branchial arch (BA) 2 directed stream displayed a polarized filopodia distribution with more filopodia from the front of the cells than the back relative to the direction of stream movement (Fig 1A). We calculated an average vector from the nucleus to the filopodia (Fig 1B) for 128 cells in the embryo shown in Fig 1A. Filopodia were defined as any thin protrusions at least 1 micron long and less than 1 micron wide extending out from the cell body. The mean number of filopodia per neural crest cell is 11.6 ± 6.6. The average filopodia vectors are strongly biased toward the direction of the migration of the overall neural crest stream (Fig 1D, p<.001), indicating that neural crest cells are morphologically polarized. Importantly, cells in the middle of the migratory stream have similar filopodial polarity as the cells at the edges of the stream (Fig 1C, Movie S1). This whole-population polarity is in contrast with the Xenopus model. Consistent with this result and in concordance with similar experiments in mice (Pryor et al., 2014), we find that PCP signaling is not necessary for neural crest cell migration in chick (Fig S1).

**Figure 1:**
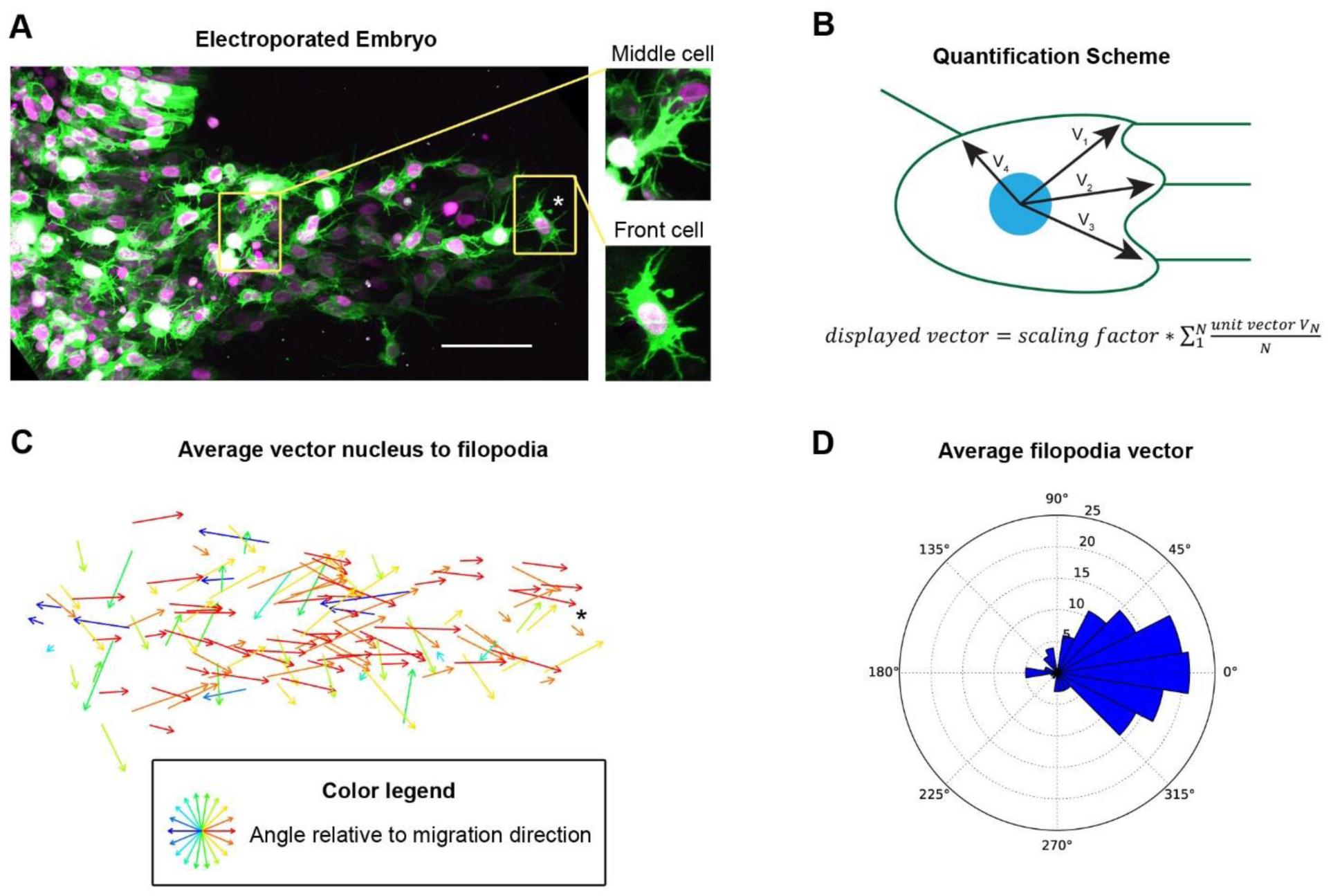
Neural crest cells throughout the migration stream are polarized. (A) HH 12 Embryo electroporated with pCAGGS-GAP43-GFP (mem-GFP, to label plasma membrane) and pCAGGS-H2B-tdTomato (to label the nucleus) and imaged live by two-photon microscopy; Image is a maximum intensity projection of 92 1 μM z-planes scale bar, 50 μM, insets are cell in middle and front of the stream. (B) Quantification scheme of average filopodia position vector. The angular position of the base of each filopodia to the nucleus relative to the direction of overall stream migration was measured in 2D. The vectors were all assigned a length of 1 and an average vector was calculated. Scaling factor is for visibility in plot. (C) Plot of vectors calculated as in panel B for 128 neural crest cells from embryo shown in panel A. Asterisk marks a cell near the front of the stream (in both 1A and 1C as a fiducial mark). Neural crest cells throughout the stream have more filopodia on their surface facing the direction of overall stream migration compared to other directions (p<.001 by Rao’s spacing test). See also Movie S1. (D) Rose plot of average filopodia position for 128 cells from embryo shown in panel A.

We next sought to determine the origin of the polarized distribution of filopodia. The biases observed in one point in time could arise from polarized generation of filopodia, polarized maintenance of filopodia, or both. To distinguish between these possibilities, we electroporated neural crest cells with mem-GFP and imaged actively migrating cells every 30 seconds for 10 minutes with a two-photon microscope to capture filopodia dynamics (Fig 2A, Movies S2 and S3).

**Figure 2:**
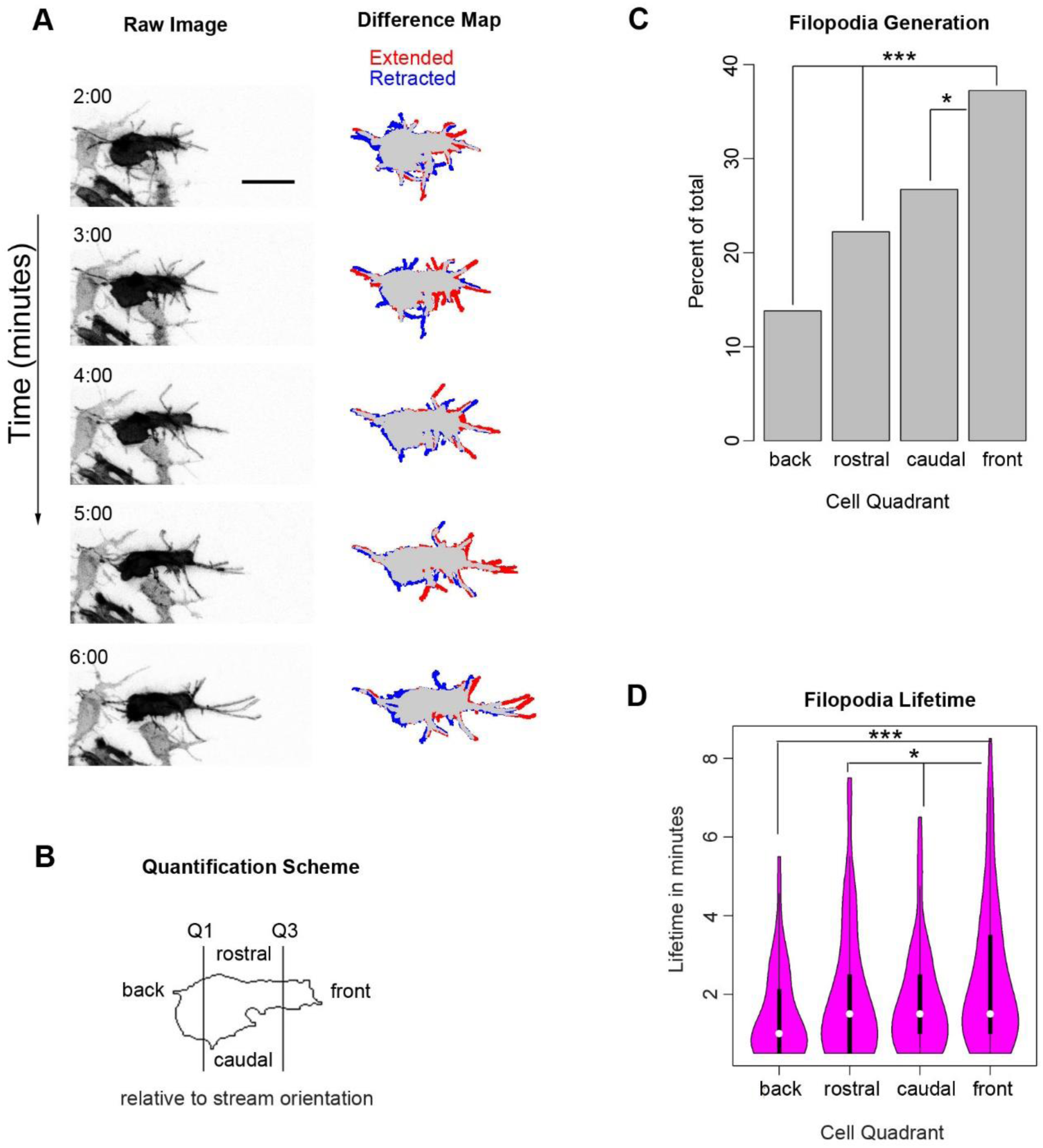
Both the generation and lifetime of neural crest filopodia are biased towards the direction of overall stream migration. (A) Still images from Movie S2 of a neural crest cell electroporated with pCAGGS-mem-GFP and imaged every 30 seconds for 10 minutes to investigate filopodial dynamics. The difference map shows a comparison of the current frame with the image taken one minute previously, with regions that were extended and retracted labeled with red and blue respectively. Time in minutes; scale bar, 20 μM. (B) Quantification scheme for filopodia spatial distribution used in graphs C and D. The perimeter of the cell body, excluding filopodia, was determined in every frame and divided into quadrants. The front and back quadrants each have 1/4 of the total perimeter while rostral and caudal split the remaining half. (C) (C and D) Filopodial generation events (C) and the lifetime of those filopodia (D) based on the quadrant where the filopodium was initiated. The direction of overall stream migration (front) has a significantly higher rate of filopodial generation and longer-lived filopodia relative to the back and sides. N=333 filopodia from 8 cells in 3 embryos. * p<.05, *** p<.001 by Chi-square or Mann-Whitney U.

Filopodia were binned according to the quadrant of the cell in which they were initiated (Fig 2B). We quantified 333 filopodia from 8 cells in 3 embryos. We find that neural crest cells have moderate biases in both filopodia generation and lifetime (Fig 2C and D). The front quadrant (corresponding to the direction of overall stream migration) has 37% of filopodia generation events, while the back quadrant has 14% of filopodia generation events (Fig 2C p<.001 by Chi-squared). The rostral and caudal quadrants even split the remaining filopodia, with 22% and 27% respectively (non-significant by Chi-squared). Filopodia lifetime exhibits a similar bias with the front quadrant having the longest lived filopodia (mean lifetime of 2.4 minutes), the back the shortest (1.5 minutes), and rostral and caudal having similar, intermediate values (2 and 1.8 minutes respectively) (Fig 2D). Thus, the robust polarity in filopodia distribution seen at an individual time point (as in Fig 2D) is a result of both biased filopodia generation as well as biased maintenance.

Filipodia are canonically considered to be sensory structures that cannot generate motility on their own (Jacquemet et al., 2015). We therefore sought to identify a connection between filopodia generation and the creation of larger-scale protrusions. We focused on neural crest cells that were generating new protrusions and quantified the generation of filopodia in the frames preceding protrusion extension. We quantified 9 protrusion generation events in 8 cells in 5 embryos. We observed a burst of local filopodia generation preceding protrusion generation by several minutes (Fig 3A and Movie S4). In the 2.5 minutes before a new protrusion was extended, new filopodia were generated in the local region where a protrusion will form at more than double the rate per unit of perimeter than in the rest of the cell (Fig 3B). This suggests a possible connection between the filopodia generation and protrusion generation.

**Figure 3:**
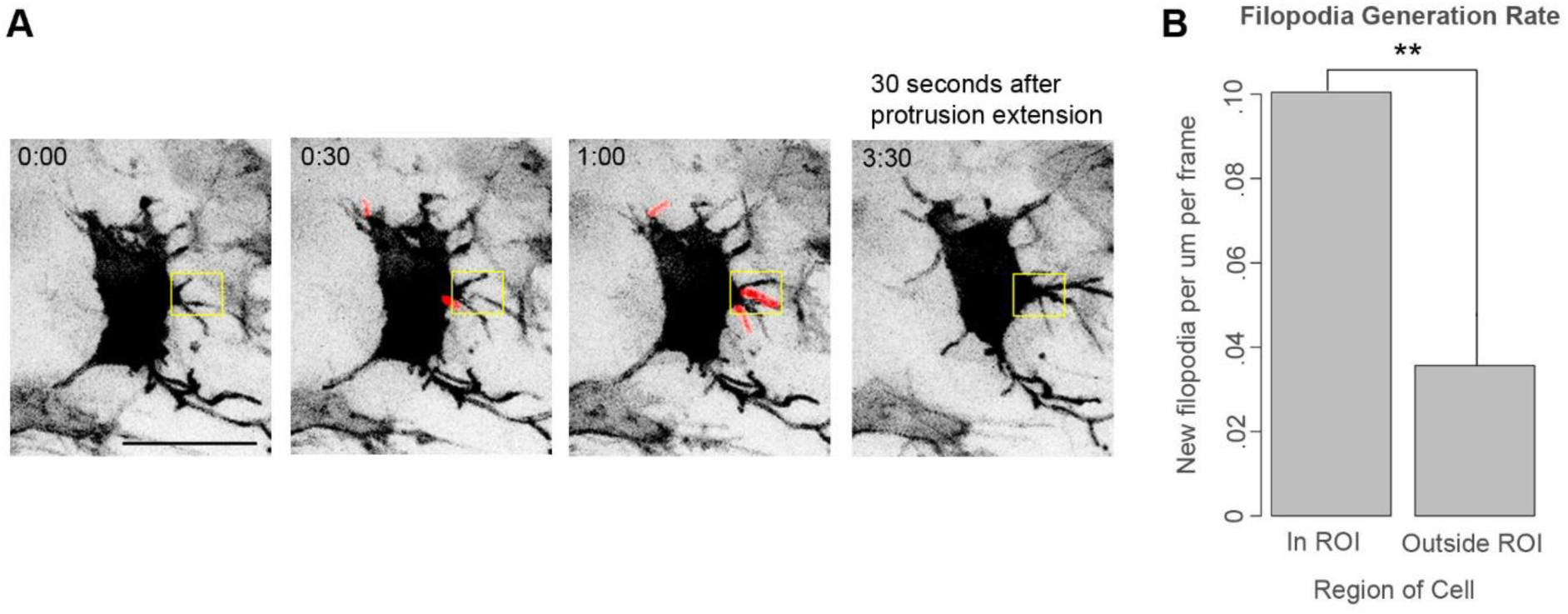
Neural crest cells locally generate a burst of filopodia before extending larger protrusions. (A) Still images from Movie S4 of a neural crest cell electroporated with pCAGGS-mem-GFP and imaged every 30 seconds. New filopodia are highlighted in red. Box is an ROI bracketing where the new protrusion will emerge. Scale bar, 20 μM. (B) Quantification of local filopodia generation rate prior to the generation of the new protrusion. Newly-generated filopodia were identified in the z-stack for 2.5 minutes prior to the extension of the protrusion. The filopodia generation rate in the ROI and the rest of the cell was calculated as a function of the cell perimeter length in those compartments. Filopodia were generated in the region where a new protrusion will emerge at more than double the expected rate compared to other areas of the cell.

Because filopodia exhibit a spatial bias toward the direction of overall stream migration and filopodia locally correlate with larger protrusion formation, we next sought to analyze the spatial bias in protrusion formation. To quantify large-scale protrusion dynamics, we sparsely electroporated neural crest with mem-GFP and H2B-tdTomato and imaged at least 90% of the neural crest stream in HH 11-12 embryos every three minutes for over two hours (Movies S5 and S6). Protrusions were defined as any structure larger than a filopodium that was actively extended from the cell body. Figure 4A shows a neural crest cell exhibiting typical behaviors such as polarizing, moving forward, and choosing between protrusions. We measured the orientation and active lifetime of 253 protrusions in 21 cells from three embryos. We defined protrusion orientation as the angle from the base of the protrusion to its tip relative to the direction of stream motion. Protrusion orientation was measured at the first frame in which the protrusion was extended. Active protrusion lifetime is the lifetime of the protrusion from birth to death excluding the retraction phase. Protrusion orientations were heavily biased toward the direction of stream movement (Fig 4B, p<.001 by Kolmogorov-Smirnov compared to a uniform distribution). They are even more polarized than filopodia generation (Fig 5C, p<.001 by Fisher exact test). Cells in the front half of the migratory stream exhibited a similar bias in protrusion orientations to those in the back half of the migratory stream (Fig S2). Protrusions oriented towards the direction of migration had a longer lifetime than those oriented anti-parallel to the direction of migration (Fig 4C, p<.01 by Mann-Whitney U).

**Figure 4:**
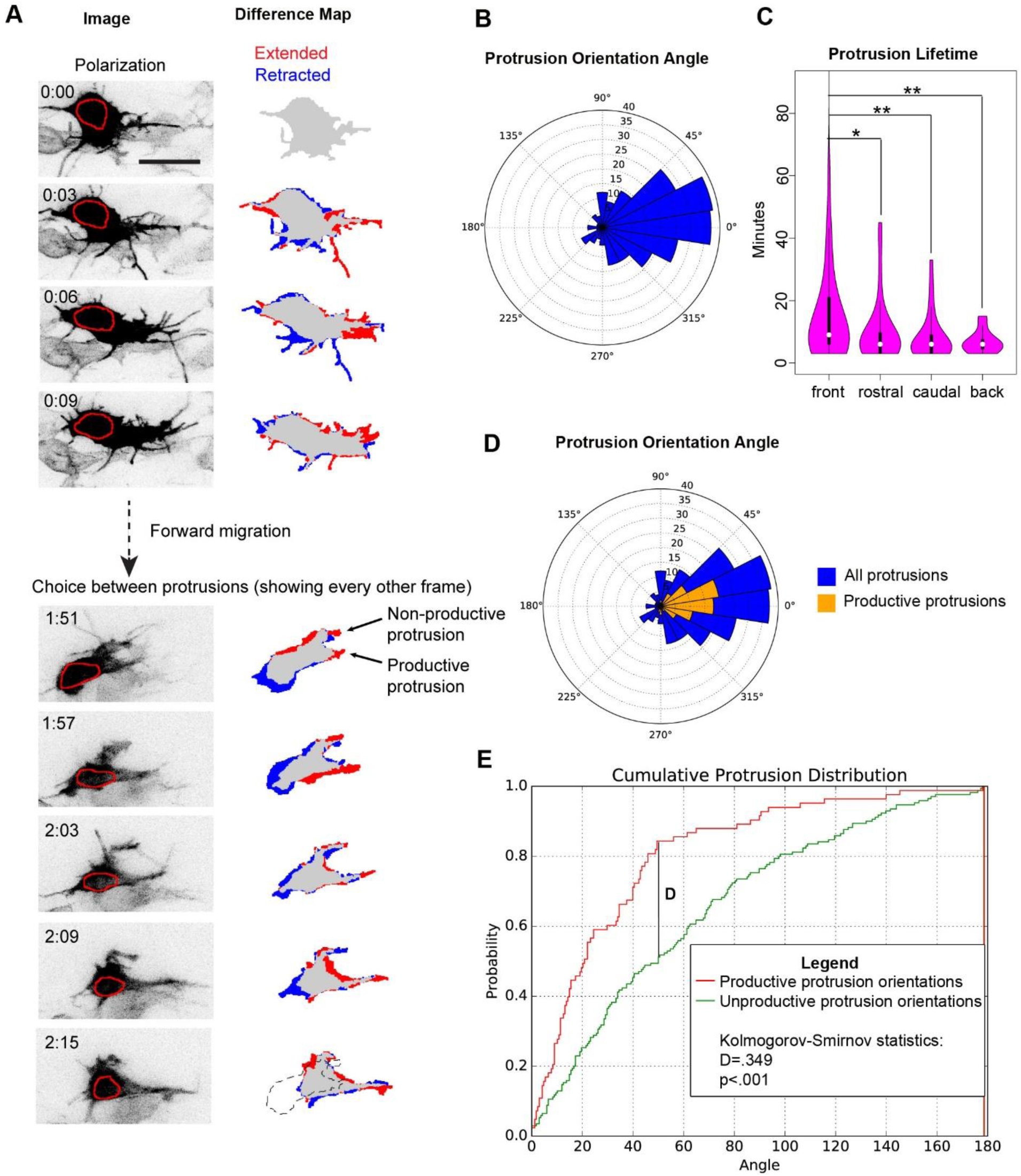
Neural crest cells migrate by biasing both protrusion generation and retraction. (A) Representative images of a neural crest cell from Movie S5 electroporated with pCAGGS-mem-GFP and pCAGGS-H2B-tdTomato and imaged every 3 minutes. The red circle outlines the nucleus. This cell exhibits typical neural crest cell behaviors such as polarization, forward migration, and choice among multiple protrusions. Difference map shows a comparison of the current frame with either the image taken three minutes earlier for the analysis of polarization (top panels) or six minutes earlier in the analysis of choice between protrusions (bottom panels), with regions that were extended and retracted labeled red and blue respectively. Dotted outline is silhouette of cell at 1:51. Time in hours:mins, scale bar 20 μM. See Movie S5 and S6. (B) Quantification of protrusion orientation angle in HH11-12 embryos. N=253 protrusions from 21 cells in 3 embryos. Protrusion angles are strongly polarized towards the direction of overall stream migration. (C) Protrusion lifetime, excluding frames when the protrusion is being retracted. Protrusions were binned by angle relative to the direction of stream movement. *p<.05, **p<.01 by Mann-Whitney U. Protrusions aligned toward the direction of overall stream movement have longer lifetimes than those protrusions that point in other directions. (D) Orientation angle of all protrusions (blue) and productive protrusions (gold). Productive protrusions were defined as ones where the nucleus enters the region that the protrusion was; i.e. what was once a protrusion is now the cell body. (E) Normalized cumulative histogram of productive and non-productive protrusion orientations. Protrusion orientations were collapsed to 0° to 180° to enable the use of more standard statistical tests. Productive protrusion orientations were more tightly clustered around the direction of overall stream movement than non-productive protrusions.

**Figure 5:**
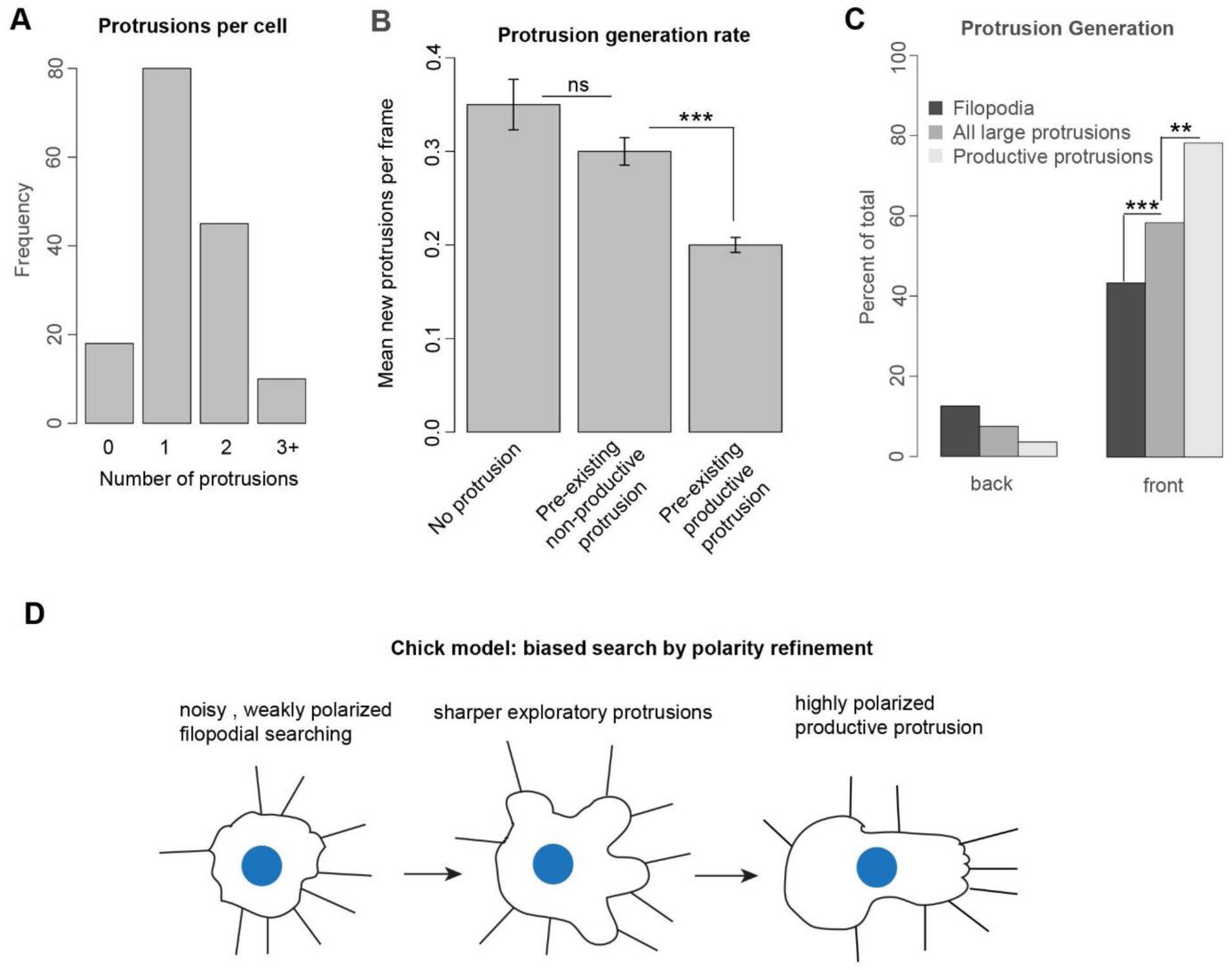
Neural crest cells exhibit competition among protrusions. (A) Number of protrusions per neural crest cell. All labeled neural crest cells, 153 in total, were quantified in a randomly chosen frame for 3 embryos. (B) Average number of protrusions generated in a 3 minute frame given the protrusive state of the cell. Neural crest cells either have no protrusions, a pre-existing not retracting *productive* protrusion, or a preexisting not retracting *non-productive* protrusion. Cells with productive protrusions have a lower rate of generating new protrusions. Error bars are Poisson s.e.m. ns is not significant. *** p<.001 by Poisson rate-ratio test. (C) Comparison of the generation rate for filopodia, all protrusions, and productive protrusions towards or away from the direction of overall stream migration. Structures are binned as in Fig 3B and 5C. There is a successive sharpening of polarization for filopodia generation, all protrusion generation, and productive protrusion generation. ** p<.01, *** p<.001, by Fisher Exact test (D) Model for chick neural crest cell migration. Chick neural crest cells migrate by refining the polarity of their protrusions.

We observed that neural crest cells frequently extend and then retract protrusions without accompanying movement of the cell body (Fig 4A). We used this behavior to classify protrusions into those that were productive versus non-productive for movement. A productive protrusion was defined as one in which the nucleus of the cell entered the previous position of the protrusion i.e. what was once a protrusion is now the cell body. Productive protrusions make up only 1/3 of the total protrusions. They have orientations that are even more tightly clustered around the direction of stream movement than total protrusions or non-productive protrusions (p<.001 by Kolmogorov-Smirnov) (Fig 4D and E, Fig 5C). These observations are consistent with a biased search model where cells efficiently explore their environment by extending multiple protrusions roughly in the desired direction before pruning the losing protrusions.

There are several possible strategies for biasing the selection of protrusions during directional migration. Cells could preferentially selecting protrusions whose individual orientation matches the desired migration direction or whose absolute position is closest to the ultimate destination (Andrew and Insall, 2007). To address this question for chick neural crest, we quantified the position relative to the center of the nucleus for every protrusion (Fig 4 and S3). Since protrusion position and orientation were reasonably correlated (Fig S3, circular Pearson’s ρ=.73), we employed a logistic regression to determine which variable was more predicative of whether a protrusion would be productive. In the linear model, only protrusion orientation had a significant coefficient suggesting that protrusion orientation is the preferred metric for evaluating protrusion quality (Fig S3).

We observed that neural crest cells frequently have multiple protrusions (Fig 4A). This was quantified by counting the number of protrusions on all labeled neural crest cells, 153 in total, in a randomly chosen video frame for three embryos (Fig 5A). Because the cell must ultimately choose one dominant protrusion to migrate efficiently, we sought to determine whether there is competition between protrusions. Specifically, does the presence of an active, productive protrusion suppress the generation of additional protrusions? To address this question, we measured the average number of additional protrusions generated per 3-minute frame while the cell has no protrusions, a non-productive protrusion, or a productive protrusion. For this analysis, the time the cell was defined as having a protrusion only included frames where the protrusion is not retracting, as we reasoned that a protrusion in the process of being dismantled had already lost the competition. We find no significant difference in the rate of new protrusion generation in a cell with no protrusions or a non-productive protrusion (Fig 5B).

However, the cells with a productive protrusion exhibit decreased generation of other protrusions (Fig 5B). This suggests that neural crest cells have a mechanism to evaluate the quality of their existing protrusions when deciding whether to extend new ones.

## Discussion

Neural crest cells exhibit progressive sharpening of the polarization of their protrusions in the direction of overall stream migration (Fig 5C). They have a moderate polarization in filopodia generation that is further sharpened by spatially biased filopodia lifetime to produce a consistently polarized final distribution of filopodia (Fig 1 and 2) aligned with the overall direction of stream migration. Bursts of local filopodia generation precede the extension of larger protrusions. Total protrusion generation is more polarized than filopodia generation, while productive protrusion orientations are the most highly polarized with 78% of productive protrusions aligned to the direction of stream migration (Fig 4E and 5C). These data suggest that chick cranial neural crest cell migration through biased search followed by polarity refinement (Fig 5D). The noisy distribution of filopodia generation allows neural crest cells to explore many possible directions for migration. The bias in filopodia lifetime further refines the region of search. The locations of high filopodial density generate larger protrusions, and a subset of the protrusions that are best aligned with the external guidance cues are used to power motility. Productive protrusions suppress the generation of new protrusions to enable processive movement (Leithner et al., 2016). This progressive polarity refinement strategy may enable neural crest cells to efficiently explore their environment and migrate accurately in the face of noisy guidance cues.

Xenopus and chick neural crest cells represent two different approaches for mesenchymal cell stream migration. In Xenopus, neural crest cells migrate as a zippered together stream in which only cells at the edge of the migratory stream are protrusive (Theveneau et al., 2010). These edge cells are then the ones that interpret guidance cues, since they principally act to modulate the lifetime of pre-existing, CIL generated protrusions (Theveneau et al., 2010). Zebrafish neural crest cells are similar to Xenopus in their requirement for PCP dependent CIL for migration, suggesting that the Xenopus model also applies to them (Matthews et al., 2008; Moore et al., 2013; Theveneau et al., 2010). In contrast, chick neural crest cells in all positions of the migratory stream are polarized and protrusive, suggesting that they all could be interpreting guidance signals. The spatial biases observed in both protrusion generation and lifetime indicate that both steps could be subject to regulation. Interestingly, mice do not need PCP signaling for neural crest migration, suggesting that their neural crest may behave more like chick (Pryor et al., 2014). Whether mouse neural crest cells have a pattern of cell polarity more similar to that of chick or Xenopus remains to be determined. Chick thus provides a model for characterizing an alternate strategy for neural crest cell migration that is still amenable to powerful live-imaging techniques. In particular, *in vivo* imaging of cytoskeletal and signaling reporters and manipulation of potential guidance cues will likely provide new insights into the molecular mechanisms of amniote neural crest cell migration.

A major open question is whether chick neural crest cells individually interpret external chemotactic cues or whether cells share information to migrate collectively. Migrating chick neural crest cells express a variety of cell communication molecules including cadherins, ephrins and Eph receptors, raising the possibility that homotypic interactions could alter neural crest cell behavior (Chalpe et al., 2010; Mellott and Burke, 2008; Nakagawa and Takeichi, 1998). However, the density of the neural crest stream combined with the complicated shapes of individual neural crest cells make it challenging to rigorously distinguish sites of interaction from non-interaction. Thus, whether contacting another cell stabilizes, destabilizes, or has no effect on protrusions remains unanswered. Perturbative experiments of candidate communication molecules will be necessary to reveal whether neural crest cells share information.

A further issue is whether neural crest cells in all regions of the migratory stream use the same migration mechanisms. Neural crest cells at the front of the stream have a different gene expression signature from those in the back (McLennan et al., 2012; McLennan et al., 2015a; McLennan et al., 2015b). Mathematical models have been developed that suggest that neural cell migration makes use of two populations of cells, leaders that directly interpret the gradient, and followers that follow the leaders (McLennan et al., 2015a; Wynn et al., 2013). We observe no difference in neural crest cell protrusive behavior between cells at the front of the stream and those at the back (Fig S2). However, this does not exclude the possibility that neural crest cells at different portions of the migratory stream are using different extracellular cues to regulate a location-invariant protrusion program. Regional inhibition of candidate molecules and pathway-specific activity reporters will be important approaches to resolve this question.

## Materials and Methods

### Embryos and Electroporations

Fertilized White Leghorn chicken eggs were obtained from Petaluma Farms (Petaluna, CA, USA) and incubated at 38° until Hamburger and Hamilton (HH) stage 8+ to 9 (Hamburger and Hamilton, 1951). A DNA solution consisting of 2.8 ug/uL total DNA (either 2.8 ug/uL of pCAGGS-GAP43-GFP or 2.3 ug/mL pCAGGS-GAP43-GFP and 0.5 ug/uL pCAGGS-H2B-tdTomato) diluted in Pannett-Compton buffer with 1 mg/mL Fast-Green was injected in the lumen of the hindbrain neural tube. Embryos were electroporated *in ovo* using a Nepagene electroporator and BTX model 516 Genetrode 1mm gold plated electrodes positioned 4.5 mm apart parallel to the embryo. Four pulses of 19 V with a 50 millisecond pulse length and a 500 millisecond pulse intervals were used.

### Sample preparation for imaging

At HH 11-12, eggs were removed from the incubator and examined. Embryos with sparse but bright labeling of the neural crest were chosen for imaging. Embryos were removed from the egg and placed in EC culture(Chapman et al., 2001). A very thin coat of EC culture was applied to a Bioptechs delta t dish (cat no. 04200417C). Embryos were sandwiched in between two rings of filter paper and clips of tungsten wire were applied to the filter rings to minimize sample drift. In order to prevent motion blur caused by the embryos’ heartbeats, the middle of the heart was carefully cut away using tungsten dissection tools 20-60 minutes before the embryo was placed on the microscope.

### Two-photon imaging

Embryos were immersed in Pannett-Compton buffer and imaged on an upright LSM 7 MP INDIMO system (Carl Zeiss Microscopy), customized with four GaAsP detectors and a Z-Deck motorized stage (Prior). Samples were imaged with a W Plan-Apochromat 20x/1.0 N.A. water-immersion objective. Excitation was with a Chameleon Ultra II laser (Coherent) tuned to 930 or 940nm, with power attenuated by an acousto-optical modulator. Images were collected with ZEN Black software. A manually adjusted Bioptechs delta t4 culture dish controller was used to maintain the embryos’ temperature between 37.5° and 38.5°. 930 nm excitation wavelength was used for samples with GFP alone and 940 nm was used for samples with GFP and tdTomato. 2x frame averaging was performed. For imaging filopodia dynamics, z-stacks with a 1 uM step size were acquired every 30 seconds for 10 minutes. For imaging protrusion dynamics, z-stacks with a 1.5 uM step size were acquired every 3 minutes for at least 2 hours. This slight under-sampling in z allowed for the whole stream to be imaged with minimal neural crest cell death and photobleaching.

### Filopodia Polarity Quantification

Filopodia were manually quantified on the z-stack using ImageJ. Filopodia were defined as long thin structures less than 1 micron thick. Nuclei XY position was measured using the Analyze Particles command on a maximum projection. Nuclei Z position was determined manually.

Vectors were calculated and the plot was generated using RStudio.

### Filopodia Dynamics Quantification

As a quality control, only bright cells with clearly visible filopodia that were still moving forward at the end of the 10 minutes were quantified. Z-stack, timelapses were opened using the FIJI distribution of ImageJ and the Correct 3D Drift plugin was applied to the raw images. Filopodia were quantified manually on the z-stack using the ROI manager. 8 cells from 3 embryos were analyzed. To aid identification of new filopodia, consecutive frames were overlaid in the green and red channels. To obtain the cell body perimeter, a maximum projection was created of the video and manually thresholded so that the entire cell body was above the threshold. Although filopodia are dimmer than the cell body, many are above the threshold. To remove the remaining filopodia and smooth the cell perimeter, the thresholded image was opened in Matlab and a morphological erosion and opening was performed. The locations of the perimeter pixels were obtained using the boundaries command. The front and back of the cell were defined as respectively, the fourth and first quartile of the perimeter along the direction of overall stream movement (Fig 2B). This ensured that those quadrants always have an equal length irrespective of cell shape. Rostral and caudal quadrants then occupied the middle of the cell. Quadrant boundaries were recalculated every frame. Filopodia were binned according to which quadrant the base of the filopodium fell into when it was first extended.

For the comparison of the location of filopodia relative to newly generated protrusions, images were processed as above to remove the filopodia. The processed image was used to define an ROI where the new protrusion was extended. These ROIs included on average 11.6% of the cell perimeter. The time the protrusion was first extended was also determined using this processed image. New filopodia were identified without reference to the ROI and then later categorized by whether the nearest perimeter pixel to the base of the filopodium was inside or outside the ROI. 9 protrusions from 8 cells in 5 embryos were analyzed.

Statistics were calculated using RStudio.

### Protrusion Quantification

Two-photon images were opened in FIJI and the Correct 3D Drift plugin was applied to the images. Cells were chosen for analysis in the first frame of the video based on brightness, separation from other labeled cells, and to ensure sampling of cells from all positions in the migratory stream. Cells that subsequently died or divided were excluded from the analysis. Occasionally during a video, a cell was cut off by stage drift or migrated into a large knot of brightly labeled cells making it difficult to discern its outline. In those cases, only frames where all protrusions could be clearly distinguished were included in the analysis. In total 21 cells from 3 embryos were analyzed.

Protrusions were quantified manually in the z-stack by creating a linear ROI along the length of the protrusion. To exclude retraction tails, protrusions were defined as any structure larger than a filopodium that was actively extended from the cell. Protrusion lifetime was measured from the first frame the protrusion was visible until it started retracting. Productive protrusions were defined as protrusions where the nuclei overlapped the ROI marking the initial protrusion location at some point in the video.

For the quantification of the number of protrusions per cell, a random frame was chosen from three videos at least 15 minutes after their start to avoid ambiguity arising from the presence of a preexisting structure, and then all labeled cells were assayed.

Statistics were calculated using RStudio.

## Acknowledgements

We thank Xin-Zi Tang and Sara Venters for helpful discussion, Rieko Asai and Lisandro Maya-Ramos for a critical reading of the manuscript and the Cardiovascular Research Institute for financial support of the two-photon.

